# The IgH *Eμ*-MAR regions promote UNG-dependent error-prone repair to optimize somatic hypermutation

**DOI:** 10.1101/2022.08.15.503996

**Authors:** Ophélie Martin, Morgane Thomas, Marie Marquet, Armand Garot, Mylène Brousse, Sébastien Bender, Claire Carrion, Jee Eun Choi, Bao Q. Vuong, Patricia J. Gearhart, Robert W. Maul, Sandrine Le Noir, Eric Pinaud

## Abstract

Two scaffold/matrix attachment regions (5’- and 3’-*MARs*_*Eμ*_) flank the intronic core enhancer (c*Eμ*) within the immunoglobulin heavy chain locus (*IgH*). Besides their conservation in mice and humans, the physiological role of *MARs*_*Eμ*_ is still unclear and their involvement in somatic hypermutation (SHM) has never been deeply evaluated. By analysing a mouse model devoid of *MARs*_*Eμ*_, we observed an inverted substitution pattern: SHM being decreased upstream from *cEμ* and increased downstream of it. Strikingly, the SHM defect induced by *MARs*_*Eμ*_-deletion was accompanied by an increase of sense transcription of the IgH V region, excluding a direct transcription-coupled effect. Interestingly, by breeding to DNA repair-deficient backgrounds, we showed that the SHM defect, observed upstream from *cEμ* in this model, was not due to a decrease in AID deamination but rather the consequence of a defect in base excision repair-associated unfaithful repair process. Our study pointed out an unexpected “fence” function of *MARs*_*Eμ*_ regions in limiting the error-prone repair machinery to the variable region of Ig gene loci.

## Introduction

The *IgH* locus, encoding the immunoglobulin heavy chain, is among the most complex in mammals, with multiple *cis*-regulatory elements controlling stepwise DNA accessibility to recombination and mutation through mechanisms that mainly rely on transcription (Perlot & Alt, 2008). Current studies of the dynamic processes that regulate chromatin conformation changes and subnuclear location have renewed interest in *cis*-regulatory regions that delimit differentially regulated chromosomal domains. Among such DNA regulatory regions, nuclear Scaffold/Matrix Attachment Regions (MARs) have been implicated in the structural and functional organization of these domains. The juxtaposition of MARs to intronic enhancer elements in both *IgH* and *IgL* loci and their conservation in humans, mice and rabbits (Scheuermann & Garrard, 1999) suggest that such regions serve physiological functions. They participate in the regulation of gene expression notably by increasing enhancer function and facilitating their action over large distances. Several proteins found to bind MARs are expressed ubiquitously or in a tissue-specific manner, respectively defining constitutive or facultative MARs (Gluch *et al*, 2008). Once attached to the nuclear matrix in a tissue specific fashion, facultative MARs could form topological barriers that could isolate or fasten chromatin regions (Gluch *et al*, 2008). Such barriers could induce DNA torsional strain with positive and negative DNA supercoiling, respectively, upstream and downstream from the RNA pol II-induced transcription bubble (Teves & Henikoff, 2014). The supercoils are then released by the action of dedicated topoisomerases (Pommier *et al*, 2016).

The *IgH Eμ* enhancer region is a combination of both the core *Eμ* (*cEμ*) enhancer element (220 bp) and two 310–350-bp flanking *MARs* (*MARs*_*Eμ*_) that were first defined by *in vitro* matrix-binding assays (Cockerill *et al*, 1987). This region, especially *cEμ*, controls early VDJ recombination events (Perlot *et al*, 2005; Afshar *et al*, 2006) and is also involved in Ig *μ* chain expression in pre-B cells (Marquet *et al*, 2014). However, its role in SHM remains unclear. An elegant model of deletion in the endogenous *Eμ* region of hybridoma cells, enforced for human AID expression, suggested the requirement of *cEμ* and a substantial function of *MARs*_*Eμ*_ for SHM (Ronai *et al*, 2005). Similarly, when added to transgenes, *cEμ* and its flanking *MARs* contribute to Ig *μ* chain expression and high levels of SHM (Bachl & Wabl, 1996; Azuma *et al*, 1993; Giusti & Manser, 1993; Motoyama *et al*, 1994; Lin *et al*, 1998; Ronai *et al*, 1999). In contrast, knock out (KO) models underlined the complexity of its physiological regulation. In a mouse model carrying the pre-rearranged VB1-8i region, *Eμ* deletion still resulted in a high level of SHM in Peyer’s patch GC B cells, arguing for a non-essential role of the enhancer (Li *et al*, 2010). More clearly, deletion of *cEμ* in the mouse germline did not reduce SHM frequency but only slightly increased the proportion of unmutated alleles; this minor effect was likely due to the reduced inflow of peripheral and, consequently, GC B cells in this model (Perlot *et al*, 2005). Strikingly, the role of *MARs*_*Eμ*_ was also elusive and somewhat controversial. Whereas their endogenous deletion, analysed in mouse chimeras by the RAG-2 complementation assay, demonstrated that *MARs*_*Eμ*_ are dispensable for VDJ recombination and IgH expression (Sakai *et al*, 1999b), the ambiguous function of *MARs*_*Eμ*_ was sustained by the discrepancy between their ability to either bind negative regulatory factors (Kohwi-Shigematsu *et al*, 1997; Wang *et al*, 1999), improve *cEμ* enhancer efficiency (Kaplan *et al*, 2001), or substitute for *cEμ* to maintain IgH expression (Wiersma *et al*, 1999). At the κ light chain locus (*Igκ*), the intronic enhancer *Eiκ* region also contains an upstream *MAR*. The implication of *MAR*_*Eiκ*_ as an enhancer of SHM was first suggested in transgenic studies (Goyenechea *et al*, 1997) and then tested in the KO mouse model that accumulated premature light chain rearrangements with a mild SHM defect (Yi *et al*, 1999), an effect comparable to one observed at *IgH* locus in hybridoma cells devoid of *MARs*_*Eμ*_ (Ronai *et al*, 2005). At that time, while these studies instigated a variety of hypotheses accounting for MARs in modulating SHM (Franklin & Blanden, 2005), these were contradicted by a study comparing *3’Eκ*-and *MAR*_Eiκ_-function in mouse KO models (Inlay *et al*, 2006). To address the controversy over the role of the scaffold in SHM, we generated a mouse model carrying a germline deletion of *MARs*_*Eμ*_ and bred it into DNA repair-deficient backgrounds. In our models devoid of *MARs*_*Eμ*_ and their *wt* counterparts, we proceeded to side by side comparison of total SHM, transcription patterns, AID targeting and error prone repair events leading to SHM, in regions located upstream and downstream from the intronic enhancer. Our study showed that the absence of *MARs*_*Eμ*_ allows some of the error-prone repair machinery to get access to the region downstream from the *Eμ* enhancer. We propose that *MARs*_*Eμ*_ act as physiological barriers for error-prone repair in activated B cells. As a rational hypothesis, our study suggests that the conservation of nuclear matrix attachment regions in Ig genes serve to optimize SHM events within the variable regions.

## Results

### Normal B cell development and Ig production in the absence of *MARs*_*Eμ*_

We generated a mouse mutant line carrying an endogenous deletion of both the 5’ and 3’ *IgH* matrix attachment regions that flank the *J*_*H*_*-C*_*H*_ intronic *cEμ* enhancer. Although generated with slightly different targeting vector backbone and homology arms, the resulting *IgH* allele, so-called *MARs*_*Eμ*_^Δ^ (Fig. 1A), is similar to that generated by Sakai et *al*. (Sakai *et al*, 1999b). Bone marrow subsets of B cell precursors were analysed in *wt* and homozygous *MARs*_*Eμ*_^Δ/Δ^ deficient mice. When compared to age-matched *wt* animals, *MARs*_*Eμ*_^Δ/Δ^ mice exhibited normal proportions and numbers of pre-proB, pro-B and pre-B cell precursors (Supplementary Table 1). Unlike endogenous deletion of the entire *Eμ* region (Marquet *et al*, 2014), *MARs*_*Eμ*_ deletion did not modify Ig *μ* heavy chain expression in early B lineage cells since proportions of IgM-expressing bone marrow B cell populations (immature, transitional and mature recirculating B cell subsets) were comparable to those of *wt* (Supplementary Table 1). Mature B cell subsets were also similar to *wt* in the spleen and peritoneal cavity of homozygous *MARs*_*Eμ*_ ^Δ/Δ^ mutants (Supplementary Table 1). In agreement with the normal inflow of mature B cells in *MARs*_*Eμ*_ ^Δ/Δ^ animals, Peyer’s patches were efficiently colonized by naive and GC B cells. Numbers and proportions of GC B cells were even significantly increased in homozygous mutants (Fig. 1B left panels and Supplementary Table 1). The similar proportion of proliferating KI67^+^ GC B cells in *wt* and *MARs*_*Eμ*_ ^Δ/Δ^ mice implied that this increase was not due to over-proliferation of Peyer’s patch B cells (Fig. 1B right panels). Finally, levels of serum Ig isotypes were unaffected in *MARs*_*Eμ*_^Δ/Δ^ animals (Fig. 1C). This normal B cell homeostasis in homozygous mutants confirmed that *MARs*_*Eμ*_ are dispensable for B cell ontogeny and antibody production. This statement is in agreement with previous studies of an analogous MAR region in the *Igκ* locus (Sakai *et al*, 1999b; Yi *et al*, 1999).

**Figure 1.**
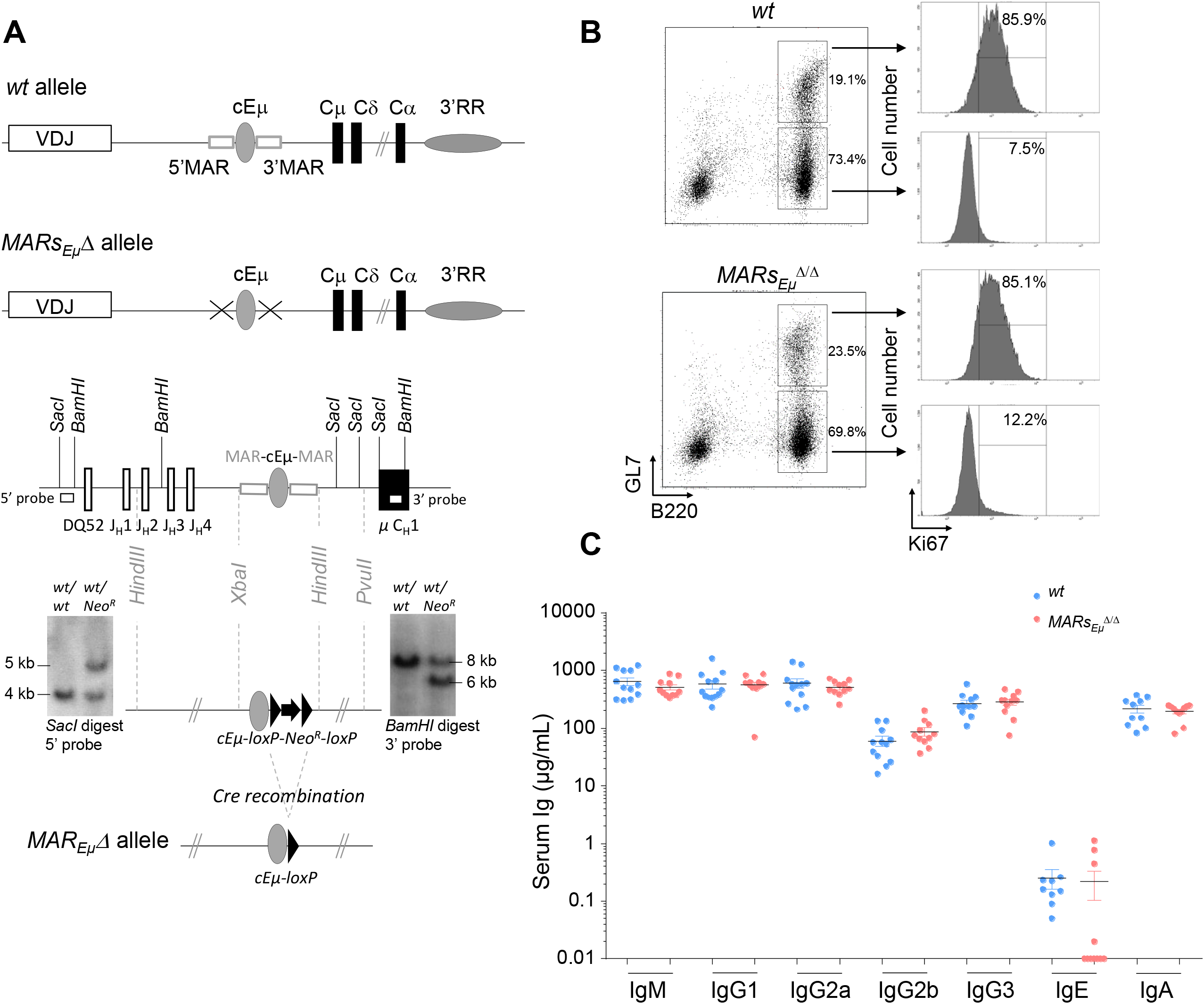
*MARs*_*Eμ*_ deletion supports efficient *in vivo* Ig isotype production and GC B cell development. **(A)** Schematic representation of *wt* and *MARs*_*Eμ*_Δ alleles (top). Targeting construct and Southern blot performed on recombinant ES cells with *Neo*^*R*^ insertion. Hybridization with the 5′ probe detected 4 kpb and 5 kpb *SacI* genomic fragments respectively for *wt* and recombined alleles. Hybridization with the 3′ probe detected 8 kpb and 6 kpb *BamHI* genomic fragments respectively for *wt* and recombined alleles. *MAR*_*Eμ*_Δ allele preserved the *cEμ* enhancer after Cre-recombination. (**B**) Comparison of Peyer’s patch B cells subsets from *wt* and *MARs*_*Eμ*_ ^Δ/Δ^ animals by flow cytometry: dot plots showed percentage of naïve (B220^+^/GL7^-^) and GC (B220^+^/GL7^+^) B cells (left panels) and, for each subset, the percentage of dividing cells (Ki67^+^) was indicated on cell count histogram plots (right panels). Experiments were performed twice with a minimum of 3 mice per group. (**C**) Immunoglobulin isotype secretion in sera from *wt* and *MARs*_*Eμ*_^Δ/Δ^ mice determined by ELISA (n=9 to 12 mice, mean±SEM).

### *MARs*_*Eμ*_ deletion inverts SHM distribution on both sides of the *Eμ* enhancer region

To assess whether *MARs*_*Eμ*_ deletion could affect *IgH* somatic hypermutation, we first quantified mutations within the 500-bp regions downstream from the variable exons rearranged to *J*_*H*_*3* and *J*_*H*_*4* segments (Fig. 2A) in Peyer’s patch GC B cells sorted from *wt* and *MARs*_*Eμ*_^Δ/Δ^ (overall data reported in Fig. 2 left, data from individual animals reported in Supplementary Figure S1A and B and Supplementary Table 2A and B). For this we used two complementary sequencing methods: the first one, based on classical Sanger approach and GS junior technology, allowed to discriminate and exclude unmutated and clonally related sequences from the calculation of SHM frequency, as initially described (Rada *et al*, 1991). The second method used Ion proton deep sequencing coupled to DeMinEr filtering. Since using DNA templates including non-mutated alleles, this second approach underestimated the SHM frequency; but since including AID-deficient control samples as a reference, the method provided highly reproducible and reliable quantification of SHM in a DNA sample extracted from GC B cells (Martin *et al*, 2018). Interestingly, by using Sanger approach, *MARs*_*Eμ*_^Δ/Δ^ GC B cells displayed significant differences in the distribution of mutations: an increased proportion of unmutated sequences (less than 10% in *wt* compared to 38% in *MARs*_*Eμ*_ ^Δ/Δ^) (from 30.6 to 45.5%, overall data collected from several mice, data from independent mice in Supplementary Fig. S1A) Another effect of *MARs*_*Eμ*_ deletion on *IgH* SHM targeting was the strong decrease in highly mutated sequences (>10bp per sequence). In *wt*, the proportion of highly mutated alleles reached∼24% (from 18.9 to 28.3%) while in mutants they were barely present (∼2%) (Fig. 2B left and Supplementary Fig S1A). When comparing only the mutated sequences, mutation frequency was decreased at least by two fold in *MARs*_*Eμ*_^Δ/Δ^ mutants, with 7.6 mutations per 1000 bp (in average) compared to 14.9 in *wt* (in average) (Fig. 2B left and Supplementary Figure S1A). By using next generation sequencing (NGS), the decreased SHM frequency was also highly significant (Fig. 2C, 12.4‰ *vs* 8.5‰, p=0.008, individual mice in Supplementary Table S2A). To monitor SHM upon antigen challenge, we analyzed mutations by Sanger and NGS in a large number of GC B cells sorted from spleen of SRBC-immunized *MARs*_*Eμ*_^Δ/Δ^ and *wt* mice. In the intronic region downstream from the *J*_*H*_*4* segment, SHM frequency dropped from 5.3 (Sanger method, excluding unmutated clones) or 3.4 (NGS bulk method) mutations per 1000bp in *wt* cells to respectively 4.6 or 2.3 in *MARs* ^Δ/Δ^_Eµ_ cells (Fig. 2B middle, and 2C, Supplementary Fig.S1B, Table S2B). Although not statistically significant, this showed that mutations accumulated by GC B cells devoid of *MARs*_*Eμ*_ were already decreased only eight days after antigen challenge. Similarly to what was observed in Peyer’s patch GC B cells, *MARs*_*Eμ*_-deficient splenic B cells also displayed an increased proportion of unmutated or poorly mutated sequences (Fig. 2B). This confirmed that the intronic region was less efficiently targeted by SHM in *MARs*_*Eμ*_ deficient mice. An identical SHM defect was also observed in mice harbouring deletion of the entire *Eμ* region (core enhancer and flanking MARs; (Marquet *et al*, 2014)) (Supplementary Fig. S1A top right); this data indicated that the SHM failure was the only consequence of *MARs* deletion. This hypothesis is completely consistent with a previous study showing that SHM efficiency was not affected by the endogenous deletion of the *cEμ* enhancer alone (Perlot *et al*, 2005). The comparison between those three models is certainly relevant since all knock outs were created in murine germlines with similar mixed genetic backgrounds.

**Figure 2.**
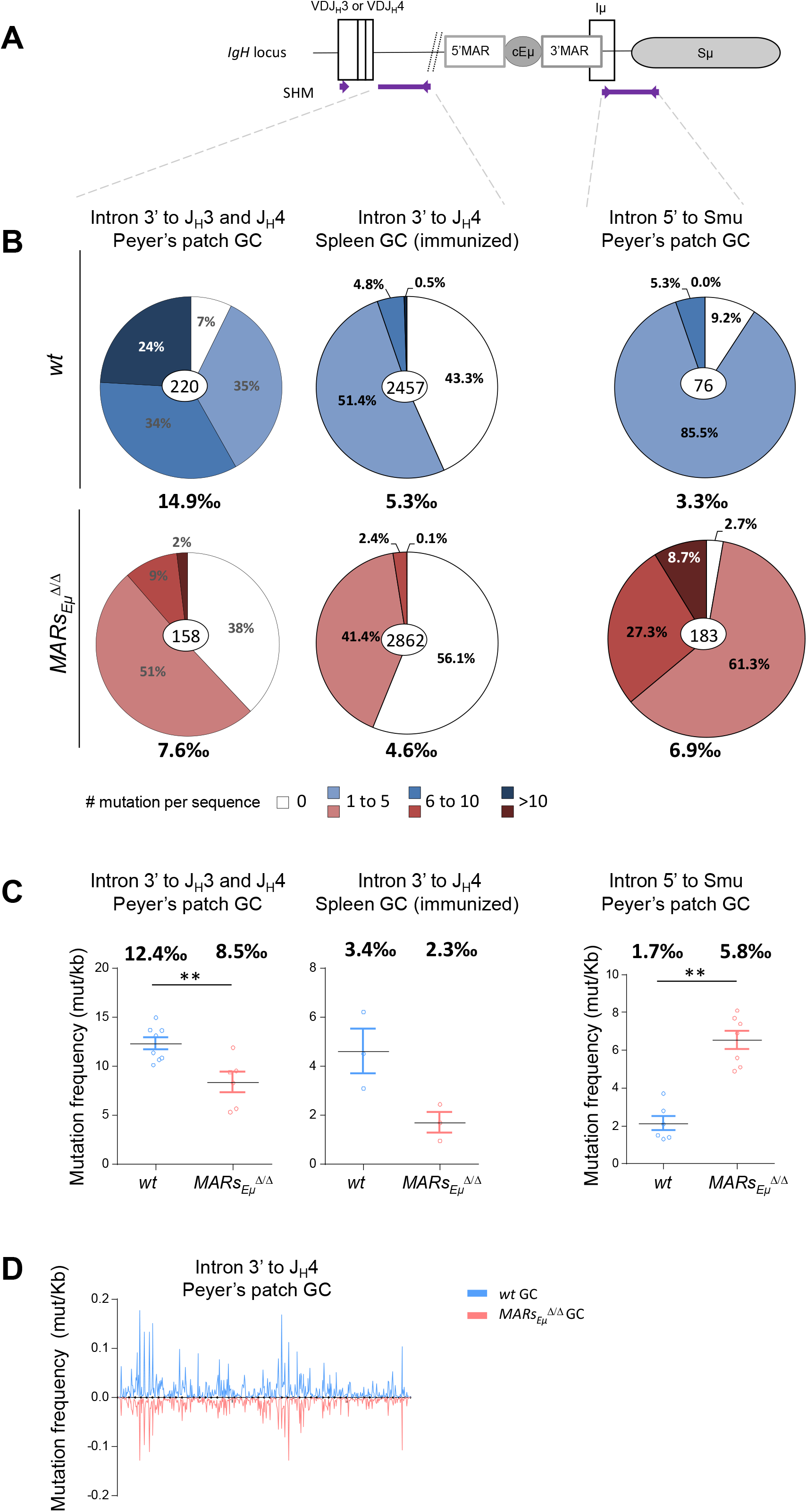
*MARs*_*Eμ*_ deletion impairs the overall SHM frequency and distribution within the *IgH J-C* intronic region. (**A**) Location of *IgH* regions (thick purple lines) tested for SHM, arrows represent primers used for PCR amplification. (**B**) Pie charts represent distribution of mutated sequences (proportional to the area in each slice, data obtained by Sanger and GS Junior sequencing method) quantified in *wt* and *MARs*_*Eμ*_^Δ/Δ^ mice in individually recombined *IgH* alleles. For each genotype number of individual clones is indicated in the center (after removal of clonally related sequences based on VDJ junction) and overall mutation frequencies (mutation per 1000 bp in mutated clones) are indicated below. Left: SHM downstream from *J*_*H*_*3* and *J*_*H*_*4* segments in Peyer’s patch sorted GC B cells, data obtained after cloning and sequencing by classical Sanger method. Middle: SHM downstream from *J*_*H*_*4* segments in spleen GC B cells sorted from SRBC-immunized mice, data obtained by NGS (GS Junior). Right: SHM downstream from *cEμ* region from Peyer’s patch GC sorted B cells, data obtained by both classical Sanger method and NGS (GS Junior). (**C**) Graphical representation of SHM frequency in *wt* and *MARs*_*Eμ*_^Δ/Δ^ mice, quantified by NGS (Ion Proton) submitted to DeMinEr filtering, a pipeline that identifies substitution frequency at each nucleotide based on an *Aicda*^*Δ/Δ*^ control sample (Martin *et al*, 2018). Since no indication in sequence distribution is available using this method, data were represented as scattered plots, each point refers to a mutation frequency from one individual mice, overall mutation frequencies are indicated above. Mean ± SEM are represented. (**D**) Mutation distribution along the *J*_*H*_*4* intron in *wt* (top) and in *MARs*_*Eμ*_^Δ/Δ^ (bottom).

Since our *MARs*_*Eμ*_ deletion includes the 3’*HinfI-XbaI* genomic region that contains transcription start sites and part of the *Iμ* exon (Lennon & Perry, 1985; Cockerill *et al*, 1987), we also quantified SHM immediately downstream from this exon in a 600bp region described as mutated in GC B cells (Fig 2A) (Nagaoka *et al*, 2002; Petersen *et al*, 2001). Unlike the intronic regions downstream from the rearranged *VDJ* exon, the overall mutation frequency downstream from *Iμ* was strongly increased in GC B cells devoid of *MARs*_*Eμ*_ region and reached 6.9 (in average) mutations per 1000 bp compared to 3.3 (in average) in *wt* cells (Fig. 2B right). This suggests that the region downstream from the *cEμ* was more efficiently targeted in the absence of its *MARs*. This was supported by the very low proportion of unmutated sequences (less than 3% in Fig. 2B right and Supplementary S1C) and the increased proportion of highly mutated sequences (more than 8%) in *MARs*_*Eμ*_-deficient GC B cells (Fig. 2B right and data from individual mice reported in Supplementary Fig. S1C). This data was efficiently confirmed by NGS analysis that estimated that mutation frequency was increased by 3.5 fold in the *MARs*_*Eμ*_-deficient GC B compared to *wt* mice (1.7‰ *vs* 5.8‰) (Fig 2C right and Supplementary Table S2C).

Analysis of mutation distribution in *wt* and in *MARs*_*Eµ*_ ^Δ/Δ^ did not show any difference between models (Fig 2D), indicating that, while affecting SHM efficiency, the absence of *MARs*_*Eμ*_ region did not influence DNA sequence hotspot or preferences for SHM within the *J*_*H*_*4* intron.

Importantly, our mouse model clearly assigns a specific function for endogenous *MARs*_*Eμ*_ on SHM at the *IgH* locus, in accord with the requirement of similar regions for efficient SHM previously pointed out in the endogenous *Igκ* Kappa light chain locus. This pioneer study, describing the specific deletion of a 420pb MAR region upstream from the intronic Kappa enhancer (*Eiκ*), highlighted a modest decrease in SHM by quantifying mutations downstream from the *J*_*κ*_*5* segment in GC B cells from Peyer’s patches (Yi *et al*, 1999). While our study suggests that *MARs*_*Eμ*_ optimizes SHM upstream from the *cEμ* enhancer; the presence of such regulatory regions does not prevent the SHM machinery to get access to downstream regions as reported in a recent study (Heltzel *et al*, 2022). This hypothesis is mostly supported by the increased SHM frequency downstream from the *cEμ* enhancer in the absence of *MARs*_*Eμ*_, a finding consistent with previous works describing increased *Sμ* internal deletions in hybridomas devoid of MARs regions (Sakai *et al*, 1999a). We could speculate that one physiological function of *MARs*_*Eμ*_ regions in GC B cells is to tightly isolate the *VDJ* transcription unit by, at least temporarily, attaching the *Eμ* region to the nuclear matrix. Such a “locked” target conformation could provide an optimal environment for somatic mutations by trapping the transcription machinery and its co-factors including AID and error-prone repair factors. This topological barrier could, at the same time, partially protect downstream constant regions from SHM; although this configuration should be brief since regions downstream from *Eμ* are also efficiently targeted by AID in GC B cells (Xue *et al*, 2006).

### *MARs*_*Eμ*_ deletion modifies transcription patterns on both sides of the *Eμ* enhancer region

It is well established that SHM in Ig *V* segments is coupled to transcription initiated at *V* promoters (Fukita *et al*, 1998). To investigate transcription-related events in SHM-targeted regions upstream and downstream from *Eμ*, we precisely quantified the total amounts of total *IgH* primary transcripts by using multiple q-PCR probes located respectively downstream from *J*_*H*_*4* and *J*_*H*_*3:* the previously described probe A (Tinguely *et al*, 2012) (Fig. 3A and Supplementary FigS2) complemented by probes A’ and C (Supplementary Figures S2 and S3A). The use of cDNA templates conducted with random hexamers showed that the amount of total IgH primary transcripts running upstream from *Eμ* did not display significant variations between *wt* and *MARs*_*Eμ*_-deficient cells in both GC and *in vitro*-stimulated samples by using probe A (Fig. 3B) as well as with probes A’ and C (Supplementary Fig. S3A), although an upward trend could be noticed in LPS-activated samples. The intriguing discrepancy between the mutation phenotype observed in *MARs*_*Eμ*_-deficient GC B cells and the silent effect on global transcription motivated a more complete study of transcription events occurring upstream from *Eμ*, particularly sense and antisense transcription since the latter has been found in cells undergoing SHM (Perlot *et al*, 2008). To proceed, we generated cDNA templates with sense transcripts, initiated at the promoter of the rearranged *VDJ* segment, with three primers located downstream from the *J*_*H*_*4* segment (S1 and S2) and within the *cEμ* enhancer (S3) (Fig. 3C and Supplementary Fig. S2 and S3A). Reciprocally, we generated cDNA templates with antisense transcripts, initiated in the intronic regions upstream from *Eμ* as described by Perlot et al. (Perlot *et al*, 2008), with four primers respectively located downstream from *J*_*H*_*2* (AS0), *J*_*H*_*3* (AS1) and *J*_*H*_*4* (AS2 and AS3) (Fig. 3E and Supplementary Fig. S2 and S3B). For both sense and antisense, quantification of transcripts was possible with the same probes A, A’ and C. For strand-specific quantification assays with a given probe, the baseline level was either provided by a control reaction (P-) measuring endogenous priming since devoid of primer or by one strand-specific template that cannot be detected by the probe (T-) as reported previously (Zhao *et al*, 2009; Bolland *et al*, 2004). To note, strand-specific transcripts were optimally detected when primers and probes were closer (sense transcripts with primer S1/probe A or antisense transcripts with primer AS2/probe A). Side by side comparison of *wt* and *MARs*_*Eμ*_-deficient activated B cells samples revealed several interesting differences. When quantified with optimal primer S1/probe A tandem, sense transcripts were significantly increased in the absence of *MARs*_*Eμ*_ (Fig. 3C). In GC B cells, a two fold increase was noticed by using S1 template (Fig. 3C, left bar graph, p=0.019). In *in vitro*-activated cells, an increase of sense transcription was also observed upon *MARs*_*Eμ*_-deletion, this effect became significant for long transcripts that reach the *cEμ* (Fig. 3C, right bar graphs, p=0.004 for S3/probe A). By using A’ and C probes, sense transcripts were hardly detectable in GC samples (Supplementary Fig. S3A; middle bar graphs); although a significant increase was noticed with S3 and S2/probe C tandems in LPS-activated samples upon *MARs*_*Eμ*_-deletion (Supplementary Fig. S3A; right bar graphs). As a potential consequence of the increased transcription of the *VDJ* unit in observed upon *MARs*_*Eμ*_ deletion, we measured by flow cytometry the level of intracellular Ig*μ* chain in Peyer’s patch naive (B220^+^/GL7^neg^) or germinal centre B cells (B220^+^/GL7^+^) of *wt* and *MARs*_*Eμ*_^Δ/Δ^ mice. In both cell types, the significant increase of intracellular Ig*μ* chain observed in the absence of *MARs*_*Eμ*_ region (Fig. 3D) corroborate our sense-transcription data. This indicated that the absence of *MARs*_*Eμ*_ certainly did not hamper RNA pol II machinery to progress 3’ to the *VDJ* unit and might even facilitate this process in activated cells.

**Figure 3.**
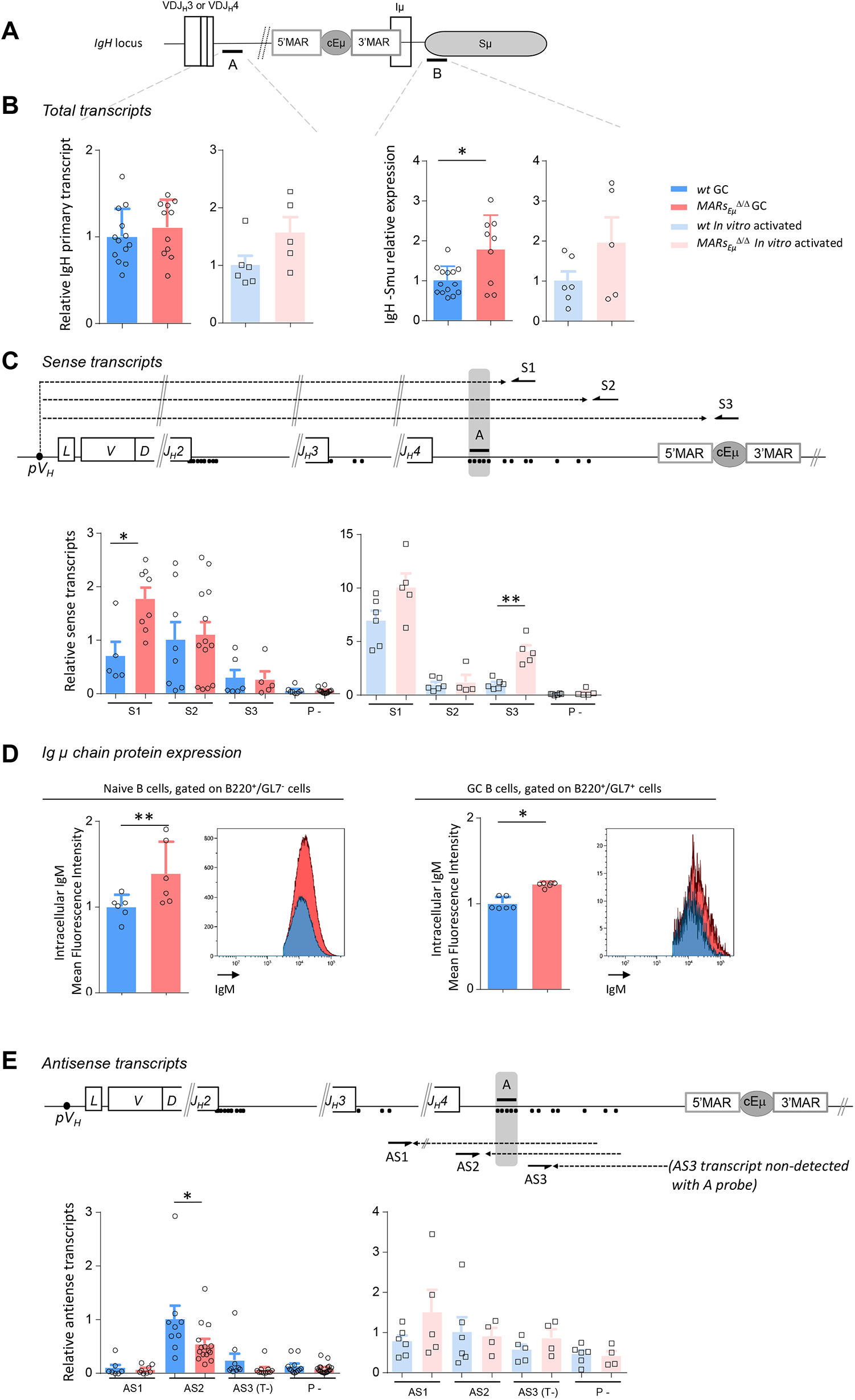
*MARs*_*Eμ*_ deletion impairs strand-specific transcription upstream from *Eμ* region. (**A**) *IgH* locus with the location of q-PCR probes (A and B) used for transcripts quantification. (**B**) Total primary transcripts quantified with A and B q-PCR probes in Peyer’s patch GC B cells (dark colors) and *in vitro*-activated B cells (light colors) from *wt* and *MARs*_*Eμ*_^Δ/Δ^ mice. (**C**) Detection of sense transcripts (dotted arrows) in murine *IgH* locus (not to scale). Arrows indicate primers (S1, S2, S3) downstream from *J*_*H*_*3 and J*_*H*_*4* used for strand-specific reverse transcription. Primary sense transcripts were quantified with A q-PCR probe (indicated by a black bar) in Peyer’s patch GC B cells and *in vitro*-activated B cells from *wt* and *MARs* ^Δ/Δ^ mice. Dots indicate antisense transcript start sites according to Perlot et *al*.(Perlot *et al*, 2008). Baseline levels were define in a control retrotranscription reaction performed without primers (P-). Bar graphs show the three (S1, S2 and S3) relative sense transcripts quantity (mean±SEM) of two to three independent experiments. (**D**) Intracellular IgM mean fluorescence intensities measured by flow cytometry in naive and GC B cells from Peyer’s patches of *wt* and *MARs*_*Eμ*_ ^Δ/Δ^ mice. Bar graphs indicate data from individual mice (*n*=6 mice in 2 independent experiments, mean±SEM); a representative example of cell count overlay is associated. (**E**) Detection of antisense transcripts (dotted arrows) in murine *IgH* locus (not to scale). Arrows indicate primers (AS1, AS2, AS3) downstream from *J*_*H*_*3 and J*_*H*_*4* used for strand-specific reverse transcription. Primary antisense transcripts were quantified with A q-PCR probe (indicated by a black bar) in Peyer’s patch GC B (dark colours) cells and *in vitro*-activated (light colors) B cells from *wt* and *MARs*_*Eμ*_ ^Δ/Δ^ mice. Dots indicate antisense transcripts start sites according to Perlot et *al*.(Perlot *et al*, 2008). Baseline levels were defined using a control retrotranscription reaction performed without primers (P-) or using a strand-specific template that cannot be detected with A q-PCR probe (T-). Bar graphs show mean±SEM of two to three independent experiments.

Globally less abundant than their sense counterparts, quantification of antisense transcripts running downstream from *J*_*H*_ segments showed quite different patterns (Fig. 3E and Supplementary Fig. S3B). While quite similar levels were detected in LPS-activated samples (Fig. 3E right and Supplementary Fig. S3B), intronic antisense transcripts were about 2 fold less abundant in *MARs*_*Eμ*_-deficient GC B cells when detection was allowed by optimal probe/primer combination (Fig. 3E, left, p=0.025, AS2/probe A and Supplementary Fig. S3B left, p=0.041 for AS3/probe A’).

The obvious unbalanced sense/antisense transcription ratio could result from either weak transcription efficiency or instability of antisense products. Nevertheless, Perlot et *al*. identified by RACE assays, in normal GC B cells, multiple antisense-transcript initiation start sites downstream from every *J*_*H*_ region and raised the question of specific enhancers. Our current data refines this previous study by identifying *MARs*_*Eμ*_ as potential boosters of antisense transcripts that, given their proximity to the enhancer, could achieve some regulatory function like eRNA or PROMPT/uaRNA (Li *et al*, 2016). Highlighting a correlation between mutation efficacy and strand-specific transcription pattern upstream from *Eμ*, our data support the idea that some level antisense transcription downstream from the *VDJ* exon could prepare to SHM (Perlot *et al*, 2008). Seemingly transient, specific to cell subsets and occurring upstream from an enhancer, such antisense transcripts could be substrates for RNA exosome and lead to optimized SHM targeting as proposed by Basu and colleagues (Lim *et al*, 2017; Laffleur *et al*, 2017, 2021).

Since a strong increase of mutations was observed within the *Sμ* region in the absence of *MARs*_*Eμ*_, we also sought to correlate SHM and transcription on the other side of *cEμ* by quantifying total transcripts (probe B) running in this region (Fig. 3A). In this case and according to what could be expected, transcription was significantly increased in *MARs*_*Eμ*_-deficient GC B cells from Peyer’s patches (Fig.3B left, p=0.04); a similar trend, although not significant, was observed in LPS activated B cells (Fig 3B left). Accordingly, we also observed a modest but reproducible increase of CSR Cγ3 and Cγ1 in *MARs*_*Eμ*_-deficient B cells stimulated *in vitro* respectively by LPS or by LPS + IL4 cocktail (Supplementary Fig. S4). A similar modest CSR effect associated to an increase of *Sμ* internal deletions has been previously reported in hybridomas carrying the same *MARs*_*Eμ*_-deletion (Sakai *et al*, 1999a). This indicated that the absence of *MARs*_*Eμ*_ lead to a global increase in transcription of the donor S region and consequently favours SHM targeting.

The significant changes in transcription patterns upstream and downstream from *cEμ* observed in our models put forward the hypothesis that *MARs*_*Eμ*_ act as physiological barriers in activated B cells, limiting sense transcription of the *VDJ* unit up to the intronic enhancer. For transcription running through the *Sμ* region, our data is in agreement with a repressive function of *MARs*_*Eμ*_ in activated B cells, in order to limit SHM targeting of this area. However, our data also suggest that *MARs*_*Eμ*_ act as transcriptional repressors of the *VDJ* unit in both naïve and activated cells; a statement in contradiction with our hypothesis that *MARs*_*Eμ*_ facilitates SHM upstream from *cEμ*. To settle such a discrepancy in our *MARs*_*Eμ*_ -deficient B cells, we first questioned AID deamination efficiency and second error-prone repair pathways processing in SHM targeted regions: upstream and downstream from *cEμ*.

### *MARs*_*Eμ*_ deletion impairs error-prone repair pathway upstream from the Eμ enhancer region

One critical experiment needed to challenge the function of *MARs*_*Eμ*_ as physiological barrier for SHM machinery was to first assess whether IgH AID targeting could be impaired in the absence of *MARs*_*Eμ*_. To proceed, we bred our *MARs*_*Eμ*_-KO mice in a genetic background deficient for both base excision repair (*Ung*^Δ/Δ^) and mismatch repair (*Msh2*^Δ/Δ^) in order to evaluate, on and unbiased manner, the DNA footprint of AID deamination upstream and downstream from *cEμ* (Fig 4 A). As expected and according to the literature (Rada *et al*, 2004; Shen *et al*, 2006; Liu *et al*, 2008), models deficient for both BER and MMR displayed only transitions at C/G pairs reflecting cytidine deamination on respectively the template and non-template strands. By looking at deamination frequencies between control (*Ung*^Δ/Δ^ *Msh2*^Δ/Δ^) and mutant animals (*Ung*^Δ/Δ^ *Msh2*^Δ/Δ^ *MARs*_*Eμ*_^Δ/Δ^), our data showed that AID activity upstream from *cEμ* was not impeded upon *MARs*_*Eμ*_-deletion; while differences were not statistically significant (evaluated on n=3 to 4 mice of each genotype), cytidine deamination even tended to be increased in B cells devoid of *MARs*_*Eμ*_, on both sides of *cEμ* (Fig 4B and Table S3A). When compared to control animals (*Ung*^Δ/Δ^ *Msh2*^Δ/Δ^), nucleotide substitution patterns were unchanged in the absence of *MARs*_*Eμ*_ (Fig S5A), proving identical strand-specific cytidine deamination: roughly 2/3 on the template strand (C to T substitutions) and 1/3 on the non-template strand (G to A substitutions). Besides imbalanced transcription upstream from *Eμ*, this data indicates that *MARs*_*Eμ*_-deletion does not impact the choice of any DNA strand for AID targeting within intronic regions.

**Figure 4.**
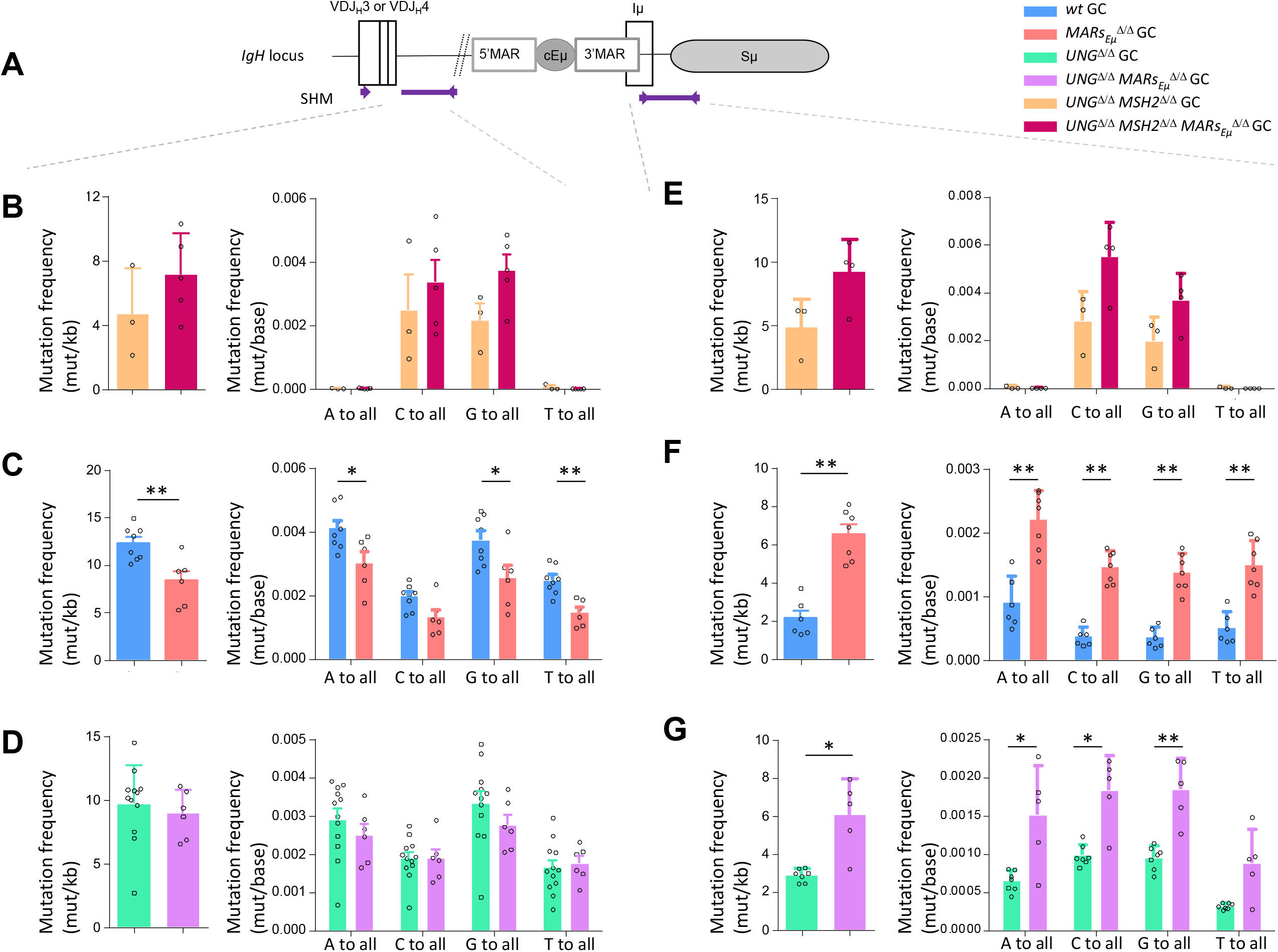
*MARs*_*Eμ*_ deletion impedes error-prone repair pathways upstream from *Eμ* region. Comparison of *IgH* SHM events occurring in Peyer’s patch GC B cells sorted from *wt* and *MARs*_*Eμ*_^Δ/Δ^ mice models, bred in genetic backgrounds deficient for base excision repair (*Ung* KO) and mismatch repair (*Msh2* KO). Data were obtained by NGS (Ion Proton) combined to DeMinEr filtering (Martin *et al*, 2018). In each region, analyzed and represented as a panel, bar graphs report overall mutation frequencies (left) and detailed mutation frequencies at all bases (right). (**A**) Location of *IgH* regions (thick purple lines) tested for SHM, arrows represent primers used for PCR amplification. (**B**) SHM downstream from *J*_*H*_*4* in double-deficient *Ung* ^*Δ/Δ*^ *Msh2*^*Δ/Δ*^ background. (**C**) SHM downstream from *J*_*H*_*4* in DNA repair proficient (*Ung* ^+/+^ *Msh2*^+/+^) background. (**D**) SHM downstream from *J*_*H*_*4* in *Ung* ^*Δ/Δ*^ background. (**E**) SHM downstream from *cEμ* in double-deficient *Ung* ^*Δ/Δ*^ *Msh2*^*Δ/Δ*^ background. (**F**) SHM downstream from *cEμ* in DNA repair proficient (*Ung* ^+/+^ *Msh2*^+/+^) background. (**G**) SHM downstream from *cEμ* in *Ung* ^*Δ/Δ*^ background. Bar graphs show mean±SEM of two to three independent experiments.

This notable increased AID deamination footprint prompted by *MARs*_*Eμ*_-deletion was in total agreement with the increased transcription observed in the corresponding regions of activated B cells. The obvious discrepancy between efficient C to U deamination events and the strong SHM targeting defect within the same *J*_*H*_ intron region unravel the origin of the SHM defect in *MARs*_*Eμ*_-deficient mice as a default of the mutagenic process occurring downstream from the normally-introduced U-G mismatches in DNA.

This prompted us to investigate whether *MARs*_*Eμ*_-deletion could provoke skewed mutation patterns within SHM-targeted regions. In the intron region downstream from *J*_*H*_*4*, mutation frequency at each of the four bases in *wt* and *MARs*_*Eμ*_-deficient backgrounds (Fig 4C and Supplementary Fig S5B) revealed a global significant decrease of mutations at all bases except for substitutions occurring at C in the absence of *MARs*_*Eμ*_. Similarly, beyond significant differences for C>A, G>C, T>A and T>G events, individual mutation patterns unveiled a global decrease that did not offer any clear hypothesis regarding the mechanism impeding SHM upon *MARs*_*Eμ*_-deletion (Supplementary Fig S5B).

To solve this paradox, we bred our *MARs*_*Eμ*_-KO mice into base excision repair deficient background (*Ung*^Δ/Δ^) and analysed SHM in the same region. Strikingly, in the absence of UNG, SHM frequency within the *J*_*H*_*4* intron region was identical upon the presence (*Ung*^Δ/Δ^ control mice) or the absence of *MARs*_*Eμ*_ (*Ung*^Δ/Δ^ *MARs*_*Eμ*_^Δ/Δ^ mice) (Fig.4D and Supplementary Table S3B). Beyond the expected increase of G/C transitions, a typical hallmark of UNG-deficient background, substitution frequencies at all four bases were also identical in *Ung*^Δ/Δ^ *MARs*_*Eμ*_^Δ/Δ^ mice (Fig 4D) the same was true when looking at individual substitution events (Supplementary Fig S5C). The fact that the SHM deficiency induced by *MARs*_*Eμ*_ deletion was no more observed in UNG-deficient background (Fig 4C and 4D) strongly imply the involvement of BER pathway in the initial mutagenic defect. This same data also proved that SHM events occurring independently of UNG (*e*.*g*. altogether obtained by replication across U and/or processed by MMR pathway) took place normally within the *J*_*H*_ intron in the absence of *MARs*_*Eμ*_. Given this, a rational hypothesis to explain the origin of the SHM defect in our model was that abasic sites generated by UNG upstream from *cEμ* are processed differently upon the absence of *MARs*_*Eμ*_. Our data suggests that U:G mismaches processed by UNG are accurately repaired in the absence of *MARs*_*Eμ*_ while these are normally subject to error-prone repair; sustaining for a specific function of *MARs*_*Eμ*_ in recruiting mutagenic BER-associated factors.

In contrast to what observed in the *J*_*H*_ intron, substitution frequencies and mutation patterns downstream from *cEμ* evidenced a different function for such regulatory regions. In B cells capable of BER and MMR, the absence of *MARs*_*Eμ*_ significantly boosted mutations at all bases by at least two fold (Fig 4F), this was true for any kind of substitution (Supplementary Fig S5E). Substitution patterns collected in mutant animals devoid of BER and MMR highlighted a global “overtargeting” of the *Sμ* region induced by *MARs*_*Eμ*_ deletion (Fig. 4E, Supplementary Fig. S5D and Supplementary Table S3C). This was in line with the general increase in both *Sμ* germline transcription observed in this model. In models impaired for BER, our data showed that UNG-deficiency combined to deletion of MARs (*Ung*^Δ/Δ^ *MARs*_*Eμ*_^Δ/Δ^ mice) maintained the SHM burden downstream from *cEμ* significantly higher than what observed for UNG alone (*Ung*^Δ/Δ^ control mice) (Fig 4G, Supplementary Fig S5F and Supplementary Table S3D). Such a comparison suggests that error-prone repair factors could more readily access to abasic sites generated in the S region when *MARs*_*Eμ*_ are missing. In this way, our data support the idea that *MARs*_*Eμ*_ act as physiological barrier that optimize SHM upstream from the *Eμ* region and rationalize the fact that MARs are evolutionary conserved downstream from Ig gene V regions (Yi *et al*, 1999); and moreover conserved structures in mammals (Scheuermann & Garrard, 1999).

### Concluding remarks

One simplistic model would argue that the most important regulatory regions for *IgH* locus expression are conserved upon any reshaping event occurring in developing B lineage cells (VDJ recombination, CSR and SHM). Beyond the major enhancer regions, *e*.*g. cEμ* and the *3’RR*, our current study identifies *MARs*_*Eμ*_, also conserved upon any rearrangement, as part of these most critical *IgH* elements. Taking advantage of the deletion of *MARs*_*Eμ*_ in the mouse, our current study rationalizes some of the molecular mechanisms that physiologically enhance SHM, through the ability of MAR to delimit some error-prone repair processes upstream from the *Eμ* intronic enhancer.

Several studies indeed proposed that *J-C* intronic MARs help generate negative supercoiling and consequently increased ssDNA and potential other secondary structures that could promote accessibility to AID (Lebecque & Gearhart, 1990; Shen & Storb, 2004; Wright *et al*, 2008). The hypothesis that *MARs*_*Eμ*_ add again more DNA strain to the sense-transcribed *VDJ* transcription unit is relevant to the positive effect of topoisomerase depletion on AID targeting and SHM (Kobayashi *et al*, 2011; Maul *et al*, 2015; Shen & Storb, 2004). A model proposed by Alt and colleagues (Meng *et al*, 2014) would be that the optimal chromatin environment for AID-induced mutations would be provided by convergent transcription as the result of fine balance between sense and antisense events. It now broadlyy admitted that RNA pol II stalling is involved in SHM-targeting (Kodgire *et al*, 2013; Maul *et al*, 2014). While DIVAC regions within Ig genes were initially described as SHM regulatory regions (Kohler *et al*, 2012), a recently study proposes that such elements facilitate RNA pol II before AID targeting (Tarsalainen *et al*, 2022). The necessity of antisense transcription for mutations is still debated, as possible byproducts of RNA Pol II collision, antisense or regulatory transcripts in such regions remain transient and difficult to detect in a *wt* context; probably because processed by RNA exosome or other RNAse activities (Basu *et al*, 2011; Pefanis *et al*, 2014). Our model clearly demonstrates that, while modifying sense and antisense transcription pattern, *MARs*_*Eμ*_ deletion does not impede AID footprint but rather some of repair mechanisms acting downstream from the U-G mismatch. Our data also indicate that only BER-dependent error-prone repair is impeded by *MARs*_*Eμ*_ deletion and suggests that efficient MMR-dependent error-prone repair in this region does not require such a barrier. The question of MARs binding factors and their respective dynamic association to such regulatory regions needs to be further investigated. The literature already suggest that some of them, like the Special AT-rich binding factor 1 (SATB1), could act as accessory factors in BER (Kaur *et al*, 2016). Recent findings showing the critical for the UNG2-interacting protein FAM72A to promote error prone processing of U-G mismatch in Ig genes (Feng *et al*, 2021; Rogier *et al*, 2021) raises the question of its specific recruitment to AID-targeted regions; our current study suggests that *MARs*_*Eμ*_ could potentially interact with UNG2 or its associated error-prone factors. Another future challenge remains to define whether some components of the nuclear matrix, nuclear filaments or proteins anchored in the envelope, could be involved in the anchorage of SHM targets.

## Material and Methods

### Generating *MARs*_*Eμ*_ KO mice

Gene targeting for matrix attachment regions flanking the *IgH Eμ* enhancer element was performed by homologous recombination, in the murine E14 ES cell line, with a vector kindly provided by Dr. Frederick Alt that permitted replacement of the 995 pb region (including *cEμ* and its flanking *MARs*) by a 220 pb *Hinf*I genomic fragment that reintroduced the *cEμ* enhancer followed by a “*loxP*-*pGK-Neo*^*R*^-*loxP*” cassette (Sakai *et al*, 1999b). Once introduced in the mouse germline, the selection cassette was deleted *in vivo* by *cre-loxP* recombination as previously described (Marquet *et al*, 2014) to obtain the *MARs*_*Eμ*_^Δ^ *IgH* allele devoid of both 5’ and 3’ *MARs*_*Eμ*_ (respectively 344 pb *Xba*I-*Hinf*I and 426bp *Hinf*I-*Xba*I genomic fragments) (Fig. 1a). Animal procedures were performed on 8 weeks old male and female mice. *wt, MARs*_*Eμ*_^*Δ/Δ*^, *Eμ*^*Δ/Δ*^ (Marquet *et al*, 2014), *Msh2*^*Δ/Δ*^, *Ung*^*Δ/Δ*^ (a kind gift of Dr S. Storck) and *Aicda*^*-/-*^ (a kind gift of Pr. T. Honjo) homozygous mice were used for our experiments and maintained at 21–23°C with a 12-h light/dark cycle. Procedures were reviewed and approved by the Ministère de l’Education Nationale de l’Enseignement Supérieur et Recherche autorisation APAFIS#16639-2018090612249522v2.

### Southern blots and PCR analysis of cre-mediated *MARs*_*Eμ*_ deletion

Genomic Southern blots were performed as follows: 20 μg genomic DNA were digested by *Sac*I or *Bam*HI and submitted to electrophoresis on a 0.7% agarose gel. DNA was transfered to nylon membranes (MP Biomedicals) by capillarity. Blots were hybridized with [^32^P]-labeled probes generated by random priming. Hybridization with 5’ probe (0.803 kpb *Sac*I-*Sph*I fragment) and 3’ probe (0.8 kpb *Xba*I-*Bam*HI fragment) located outside the homology arms were used to identify ES cell clones in which *MARs*_*Eμ*_ were replaced by the *loxP*-*pGK-Neo*^*R*^-*loxP* cassette (Fig. 1a).

### Total serum Ig quantification by ELISA

Sera were collected at 8 weeks of age from non-immunized *wt* and *MARs*_*Eμ*_^*Δ/Δ*^ mice and analyzed for the presence of different Ig classes and subclasses by ELISA as previously described (Pinaud *et al*, 2001).

### SRBC Immunisation

Mice were challenged by intraperitoneal injection with 200μL 50% sheep red blood cell suspension and sacrificed 8 days later to collect GC B cells in the spleen.

### Flow cytometry and cell sorting

Flow cytometry analysis was performed on LSR-Fortessa cell analyzer (BD Biosciences) on single-cell suspensions from fresh organs. Once washed with 2% fetal calf serum-PBS, lymphoid cells from bone marrow, spleen, peritoneal cavity and Peyer’s patches were labeled with various conjugated Abs: αB220-V450, αCD117-PE, αCD43-PE for bone marrow cells. αB220-V450, αCD21-PE, αCD23-FITC, αIgM total-PE, αIgD-FITC and αCD3e-FITC for splenocytes. αB220-V450, αIgM total-PE, αCD5-FITC for peritoneal cavity. αB220-V450, αB220-APC, αIgA-FITC, αIgM total-PE, αPNA-FITC, αFAS-PE, αKi67-FITC for Peyer’s Patches. (Southern Biotechnology Associates; eBioscience; Sigma and BD Biosciences). Flow cytometry cell sorting was performed on an ARIA 3 (BD Biosciences) apparatus on single-cell suspensions from spleens or Peyer’s patches. Once washed with 2% fetal calf serum-PBS, cells were labeled with PNA, GL7, αB220, and αFAS reagents and sorted based on distinct gates defined as germinal centre B cells (B220^+^/GL7^+^ or B220^+^/PNA^High^/Fas^+^).

### Cell culture

Splenocytes were collected, after red blood cells lysis, CD43^+^ cells were depleted using anti-CD43 MicroBeads (Miltenyi Biotec). CD43^-^ splenic B cells were cultured for 3 days at a density of 1×10^6^ cells per mL in RPMI 1640 supplemented in 10% serum calf fetal, sodium pyruvate (Lonza), amino acid (NEAA 100x Lonza) and Penicillin-Streptomycin (Gibco) with 1μg/ml LPS (Invivogen) alone (for transcription assays) or plus 20ng/ml IL4 (PeproTech) (for CSR experiments).

### SHM assays

SHM analysis was either performed by cloning followed by classical Sanger method as described (Rouaud *et al*, 2013) or performed directly on PCR products by next generation sequencing using GS Junior (Roche) or Ion Proton system (Applied Biosystem). For GS Junior, sequencing libraries were prepared according to the manufacturer’s instructions, adaptor sequences were added to the previous amplification primer sequences in order to be compatible with the GS-Junior sequencing technology. Amplifications were performed with Phusion® High-Fidelity DNA Polymerase (New England Biolabs) according to the following program: DNA was denatured 40 s at 98°C and then submitted to 38 cycles consisting of 98°C for 10 s, 68°C for 30 s and 72°C for 30 s, and 1 cycle at 72°C for 10 min. PCR products were first purified using NucleoSpin kit (Macherey-Nagel) followed by Ampure bead purification (Beckman Coulter). PCR products were subjected to “PCR emulsion step” (GS Junior+ emPCR Kit (Lib-A), Roche) and sequenced using GS Junior sequencing kit XL+ (Roche) according to the manufacturer’s instructions. Raw sequences were aligned against reference sequences of *IgHJ*_*4*_-downstream intron or *Smu* and only full length sequences were kept for mutation analysis. For *IgHJ*_*4*_, clonally related sequences were removed based on the sequence of VDJ junction similarity. No further filtering steps were implemented in our analysis workflow. Mutations were called on each sequence using pairwise alignment algorithm (from biopython package) and only base substitutions were reported. Mutation frequencies were computed as the ratio between the sum of mutated bases in all complete sequences over the total number of aligned bases. For Ion Proton, sequencing libraries were prepared according to the user guide Ion Xpress**™** Plus gDNA Fragment Library Preparation (Cat. no. 4471269, Life Technologies). Briefly, PCR products (100ng) were fragmented by enzymatic digestion (Ion Shear**™** Plus Reagents Kit, Cat. no. 4471248) and ligated to Barcodes and Adapters (Ion Plus Fragment Library Kit, Cat. no. 4471252). After 200 bp size selection step on E-Gel precast agarose electrophoresis system, final amplification was performed. Raw data were processed using DeMinEr tool as described (Martin *et al*, 2018).

### RT-PCR and q-PCR

Total RNA was prepared by using TRIzol reagent (Ambion) procedures. RNA samples were first treated with DNase I (Invitrogen) for 15 min at 25°C. RT was performed on 200 ng of total RNA with random hexamers or with specific primer (10μM) (sequence available in Supplementary Fig. S2.) using superscript III enzyme (Invitrogen). As control, we performed a reverse transcription without primer to determine the threshold (referred ad P^-^ in bar graphs). Each real-time qPCR reaction was performed in duplicate on 10 ng of RNA equivalent, using TaqMan Universal (except for Sμ quantification we used SYBR green Mastermix (TAKARA)) on StepOnePlus system (Applied Biosystems). Primary transcription at *IgH* locus was quantified as previously described (Tinguely *et al*, 2012) and completed with both a set of primers and q-PCR probes close to *J*_*H*_ segments (listed in Supplementary Fig.2.) and a set of primers located 5’ to *Sμ* : Smu-Fw (5’-ACCCAGGCTAAGAAGGCAAT-3’), Smu-Rev (5’-CCTTCCTTCTGCGTATCCAT-3’). Relative mRNA levels were normalized to *Gapdh* transcripts with the appropriate TaqMan probe (Mm99999915_g1, Applied Biosystem). Data were analyzed by comparing threshold cycle (*CT*) values according to the 2^-(ΔΔ*CT*)^ method. The *wt* mice templates used as calibrators were S2 for sense transcripts, AS0 or AS2 for antisense transcripts.

### Statistical analysis

If not specified in the figure legend, Mann Whitney two-tailed tests were used for statistical analysis using GraphPad Prism software (*p<0.05, **p<0.01, ***p<0.001, ****p<0.0001).

### Data availability

NGS data are available at the European Nucleotide Archive (PRJEB52221).

## Acknowledgments

We thank BISCEm and the animal core facility team for help with mouse work on both practical and regulatory aspects. We are grateful to Drs. Fred Alt for providing *MARs*_*Eμ*_ targeting constructand Zéliha Oruc-Ratinaud regarding the animal health status. We thank Drs. Dominik Schenten, Sébastien Storck, Christophe Sirac, Laurent Delpy, Brice Laffleur, Alexis Saintamand and Jeanne Moreau for discussions and helpful comments. OM and MT were supported by PhD fellowship of the french Ministère de l’Enseignement Supérieur, de la Recherche et de de l’Innovation. This work was supported by Région Nouvelle Aquitaine, La Ligue Contre le Cancer (comités 87, 23 to EP and SLN); the Fondation ARC pour la recherche sur le cancer (PJA 20181207918 to EP and PhD continuation fellowship to MT), Institut CARNOT CALYM, INCa-Cancéropôle GSO Emergence (to EP).

## Author contribution

OM, MT, MM, AG, MB, SB, CC, JEC, EP and SLN performed experiments. EP and SLN conceived and supervised the study. MM developed the experimental model. OM, BQV, PJG, RWM, EP and SLN wrote the manuscript.

## Figure legends

**Supplementary Figure S1.**
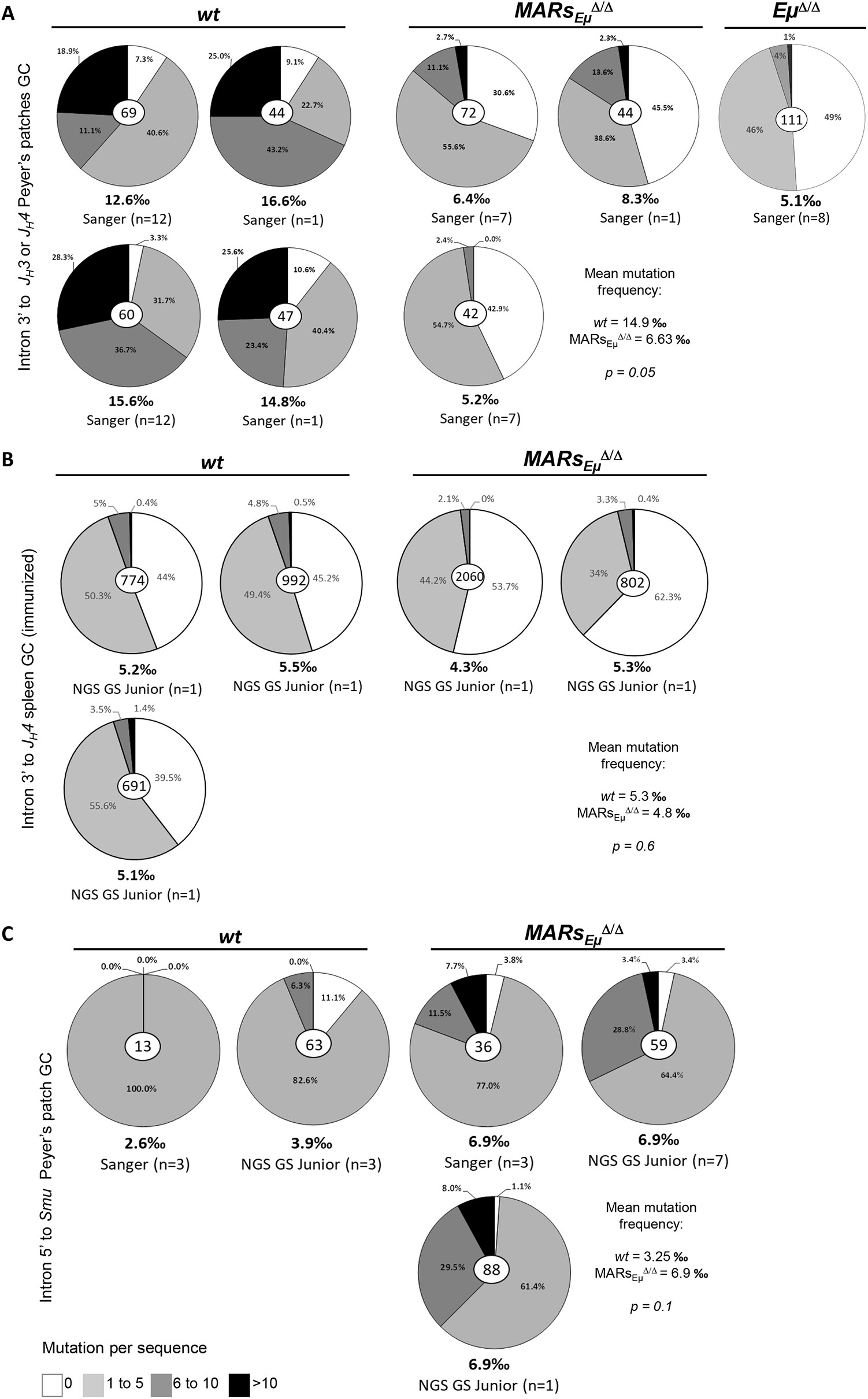
*MARs*_*Eμ*_ deletion inverts SHM distribution on both sides of the *Eμ* enhancer region. (**A**) SHM downstream from *J*_*H*_*3* and *J*_*H*_*4* segments in Peyer’s patch GC B cells sorted from *wt* and *MARs*_*Eμ*_^Δ/Δ^ mice; after cloning and sequencing by classical Sanger method. For each genotype, pie charts represent distribution of mutated sequences (proportional to the area in each slice, data obtained by Sanger and GS Junior sequencing method) in individually recombined *IgH* alleles. Number of individual clones is reported in the center (after removal of clonally related sequences based on VDJ junction). Each pie chart represent SHM obtained from an individual experiment. Under each pie chart, SHM frequency, sequencing strategy and sample type (individual mice or pool) is indicated. Mean SHM frequency and p values are reported. (**B**) Equivalent data representation than reported in **A** for SHM downstream from *J*_*H*_*4* segments in splenic GC B cells sorted from SRBC-immunized *wt* and *MARs*_*Eμ*_^Δ/Δ^ mice. Mean SHM frequency and p values are reported. (**C**) Equivalent data representation than reported in (**A**) for SHM within the intron 5’ to *Sμ* region in Peyer’s patch GC B cells sorted from *wt* and *MARs*_*Eμ*_^Δ/Δ^ mice

**Supplementary Figure S2.**
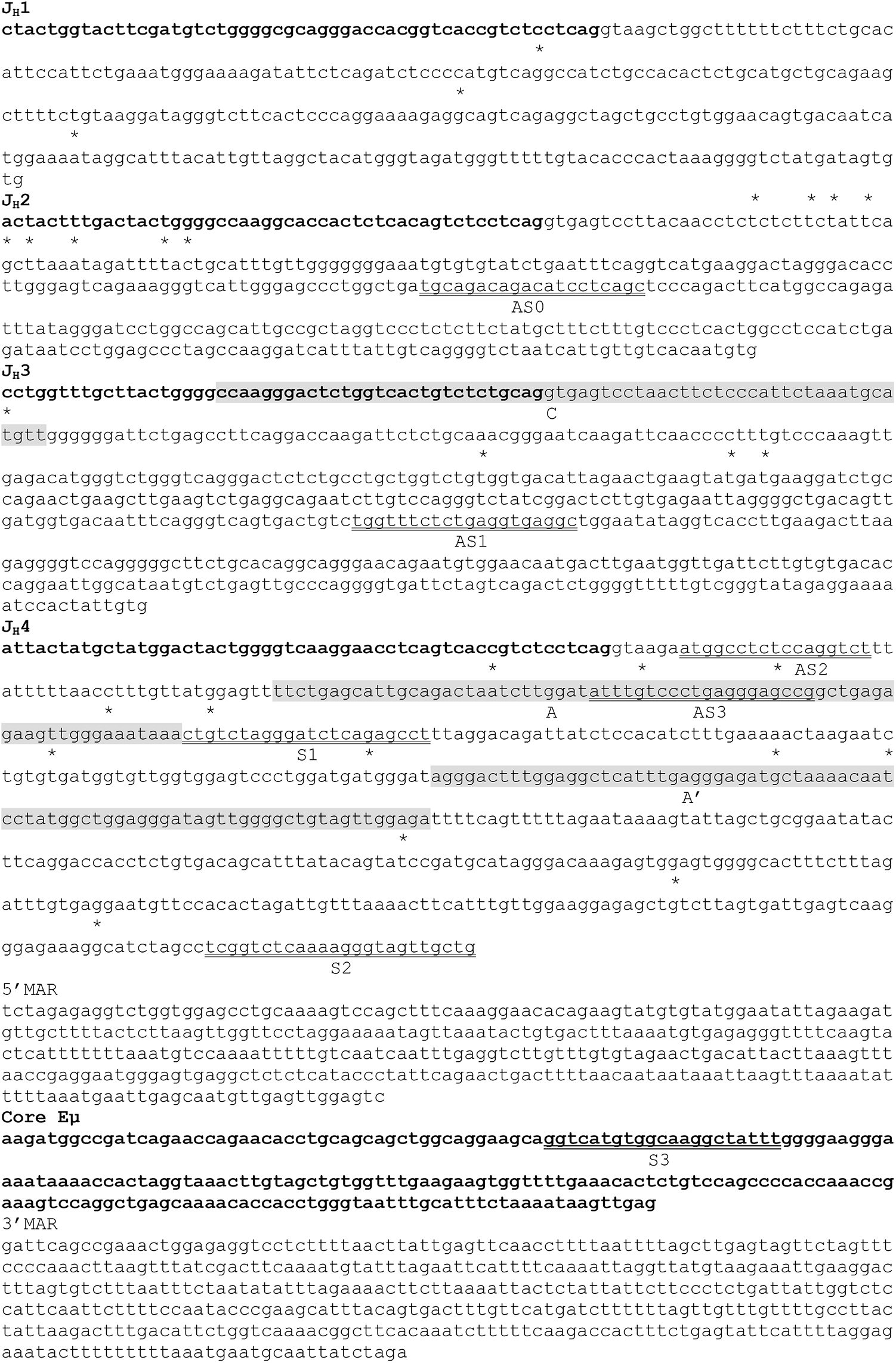
Annotated nucleotide map of the *IgH*-*J*_*H*_*1* to *Eμ* germline region from of 129 wt mice. All *J*_*H*_ exons as well as *coreEμ* element are indicated by bold characters. Start sites for antisense transcripts are reported as (*) according to Perlot et *al*.(Perlot *et al*, 2008). Location of primers used for strand-specific reverse transcription (S1, S2, S3, AS0, AS1, AS2, AS3) are indicated by underlines. TaqMan qPCR amplicons (C, A, A’) are highlighted in grey.

**Supplementary Figure S3.**
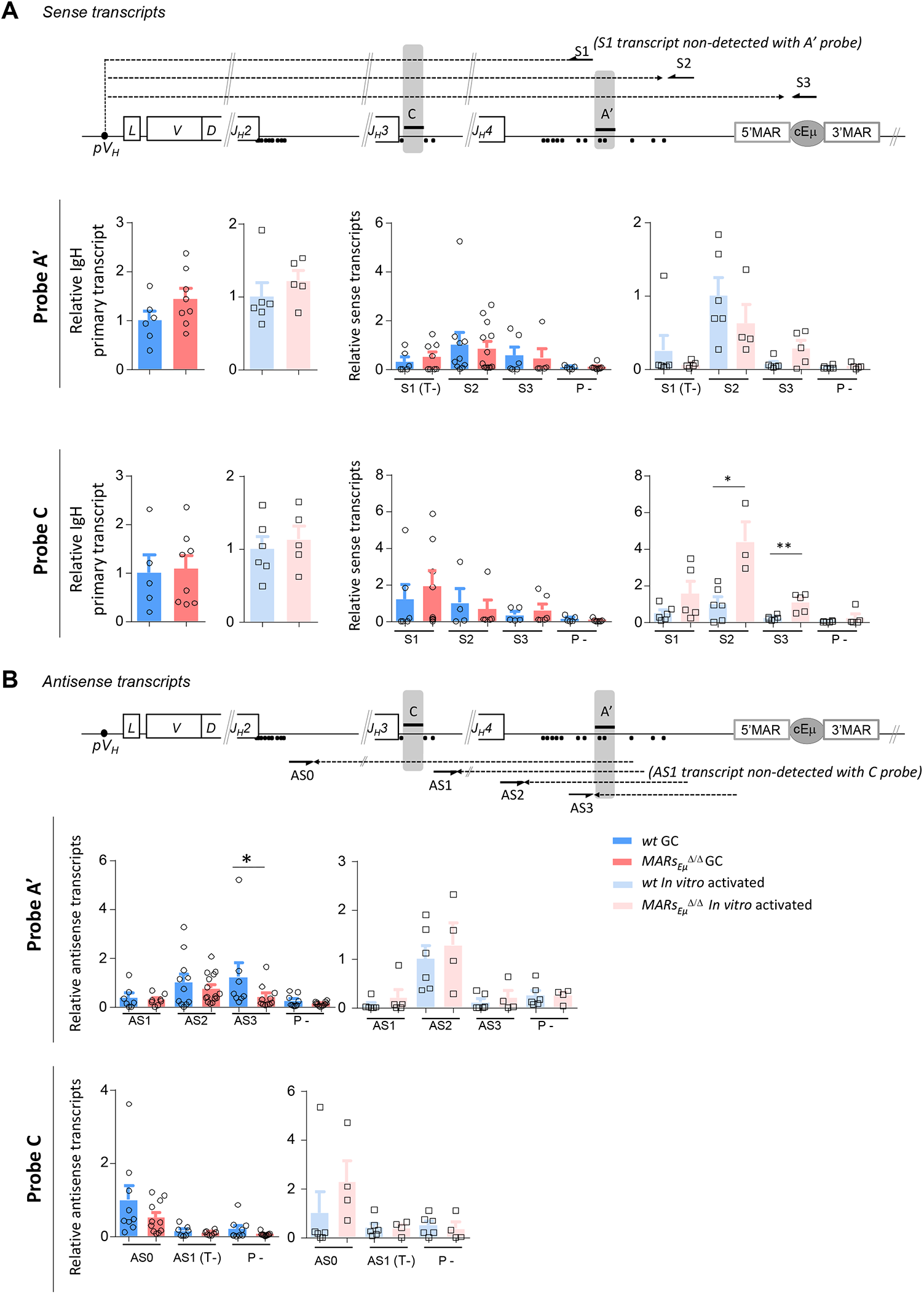
Sense and antisense transcripts quantified with *IgH J*_*H*_*3* and *J*_*H*_ *4* exons with additional TaqMan probes. Murine *IgH* locus (not to scale) indicating location of primers (S1, S2, S3; black arrows) within introns downstream from *J*_*H*_*3 and J*_*H*_*4* used for strand-specific reverse transcription to detect sense transcripts (dotted arrows). Black bars (A’ and C) indicate location of q-PCR probes. Total primary transcripts and primary sense transcripts were quantified with A’ and C probes in Peyer’s patch GC B cells (filled bar graphs) and *in vitro*-activated B cells (bar graphs) from *wt* and *MARs*_*Eμ*_^Δ/Δ^ mice. Dots indicated antisense transcripts start sites according to Perlot et *al*.(Perlot *et al*, 2008). (**B**) Murine *IgH* locus (not to scale) indicating location of primers (AS0, AS1, AS2, AS3; black arrows) within introns downstream from *J*_*H*_*2, J*_*H*_*3 and J*_*H*_*4* used for strand-specific reverse transcription to detect antisense transcripts (dotted arrows). Black bars (A’ and C) indicate location of q-PCR probes. Primary antisense transcripts quantified with A’ and C probes in Peyer’s patch GC B cells (filled bar graphs) and *in vitro*-activated B cells (emptied bar praphs) from *wt* and *MARs*_*Eμ*_ ^Δ/Δ^ mice. Dots indicated antisense transcripts start sites according to publised data (Perlot *et al*, 2008). Baseline level was either provided using a control retrotransciption reaction performed without primers (P-) or using one strand-specific template that cannot be detected with the current probe (T-). Bar graphs show mean±SEM of two to three independent experiments.

**Supplementary Figure S4.**
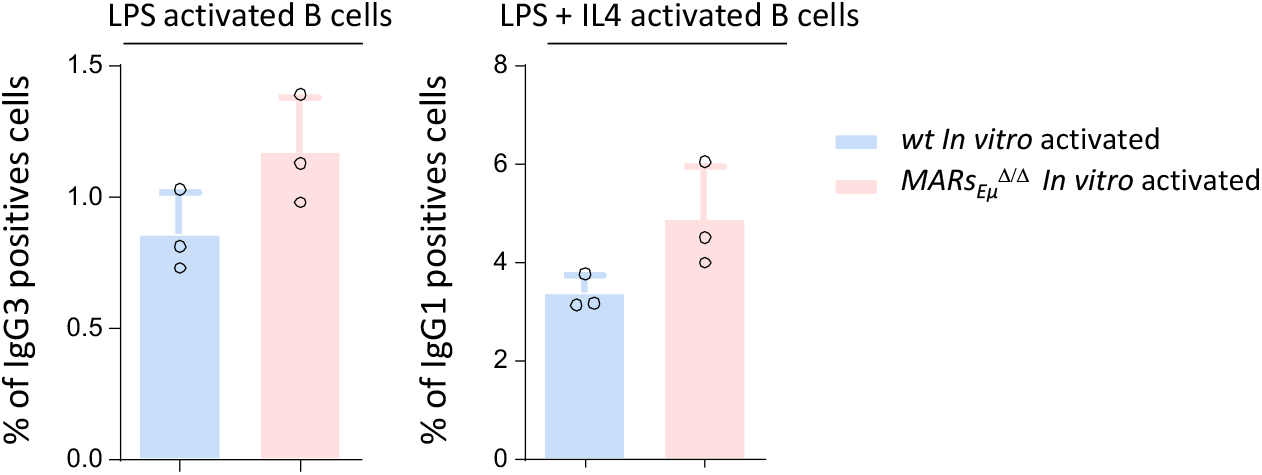
Comparison of *in-vitro* Ig class switching in *wt* and *MARs*_*Eμ*_^*Δ/Δ*^ mice. Percentage of IgG3 and IgG1 positive cells measured by flow cytometry after respectively LPS or LPS + IL4 stimulation for 3 days of splenic B cells sorted from *wt* and *MARs*_*Eμ*_ ^Δ/Δ^ mice.

**Supplementary Figure S5.**
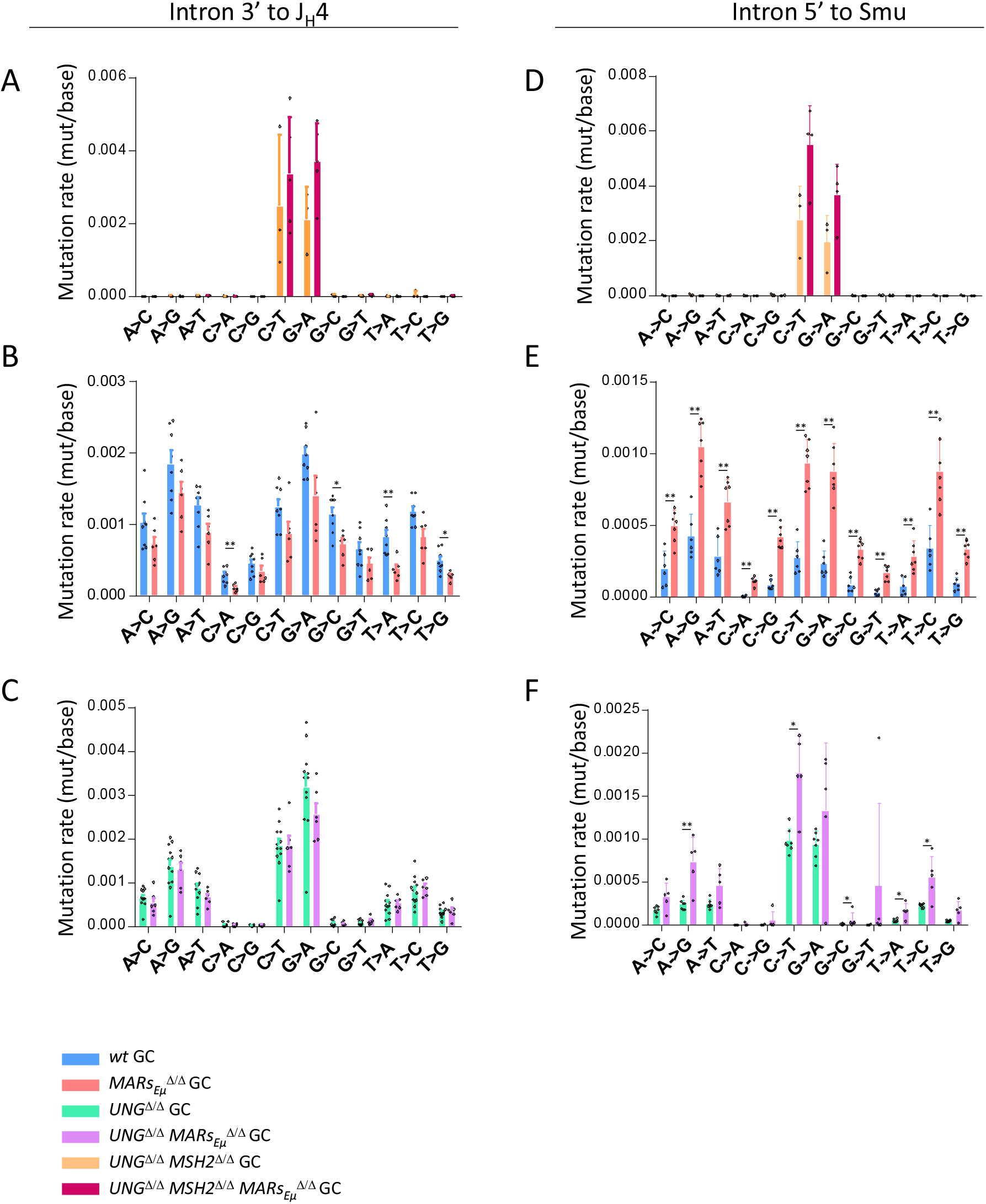
Base substitution patterns in BER- and MMR-deficient backgrounds. Comparison of SHM-related base substitution patterns, reported as frequencies, at *IgH* in Peyer’s patch GC B cells sorted from *wt* and *MARs*_*Eμ*_^Δ/Δ^ mice models, bred in genetic backgrounds deficient for base excision repair (*Ung* KO) and mismatch repair (*Msh2* KO). Data were obtained by NGS (Ion Proton) combined to DeMinEr filtering (Martin *et al*, 2018). (**A**) Substitution pattern downstream from *J*_*H*_*4* in double-deficient *Ung*^Δ/Δ^ *Msh2*^Δ/Δ^ background. (**B**) Substitution pattern downstream from *J*_*H*_*4* in DNA repair proficient (*Ung*^Δ/Δ^ *Msh2*^Δ/Δ^) background. (**C**) Substitution pattern downstream from *J*_*H*_*4* in *Ung*^Δ/Δ^ background. (**D**) Substitution pattern downstream from *cEμ* in double-deficient *Ung*^Δ/Δ^ *Msh2*^Δ/Δ^ background. (**E**) Substitution pattern downstream from *cEμ* in DNA repair proficient (*Ung*^Δ/Δ^ *Msh2*^Δ/Δ^) background. (**F**) Substitution pattern downstream from *cEμ* in *Ung*^Δ/Δ^ background. Bar graph show mean±SEM of two to three independent experiments.

**Table S1.**
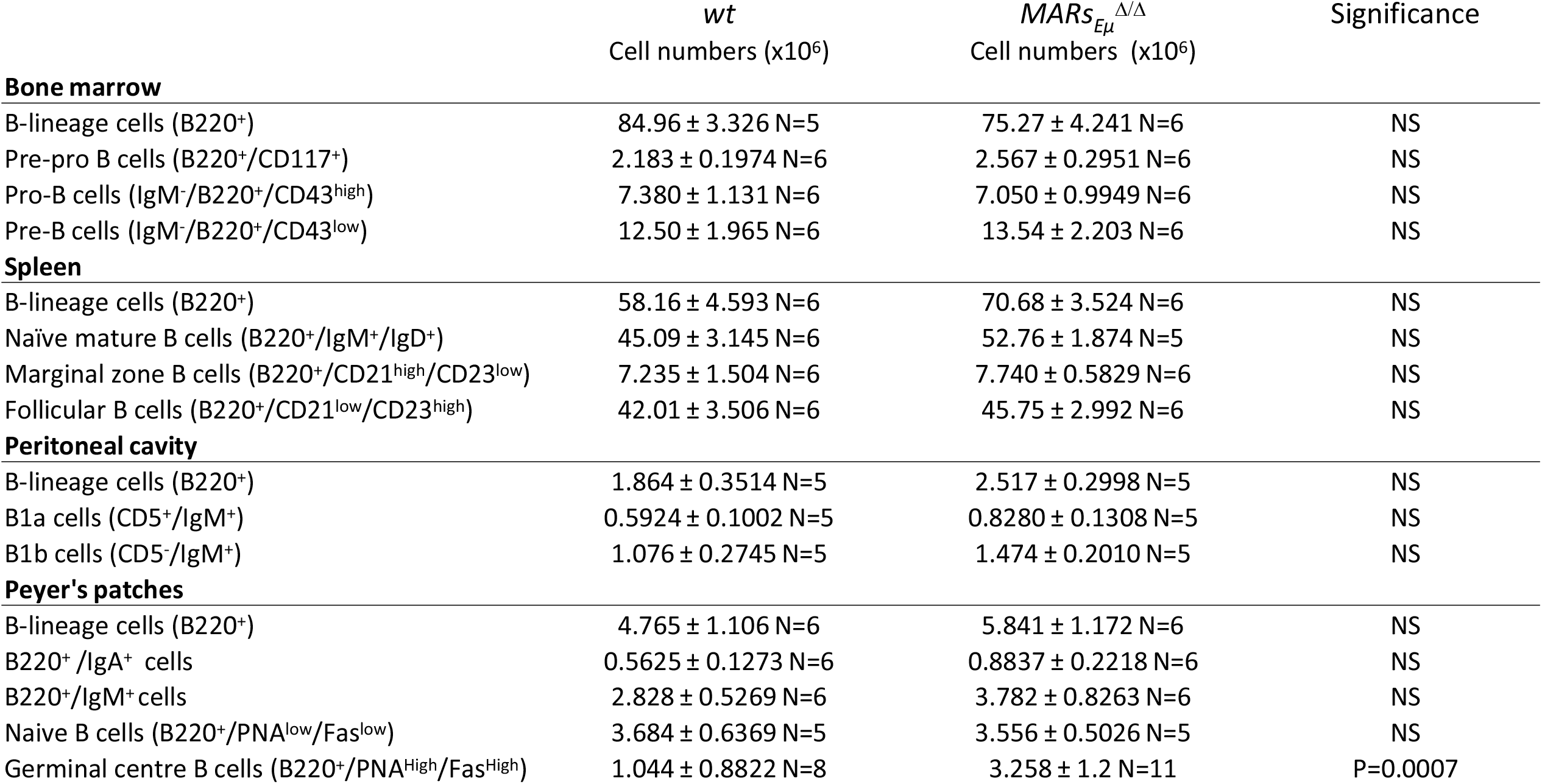
*MARs*_*Eμ*_ deletion led to normal B-lineage cell development. Bone Marrow and peripheral B cell subsets counts in *wt* and *MARs*_*Eμ*_^Δ/Δ^ mice. Absolute numbers are reported as mean± SEM. Significance was assessed with Student T test. P value is indicated when difference is significant.

|  | *wt*<br>Cell numbers (x10<sup>6</sup>) | *MARs*<sub>Eμ</sub><sup>Δ/Δ</sup><br>Cell numbers (x10<sup>6</sup>) | Significance |
| --- | --- | --- | --- |
| **Bone marrow** |  |  |  |
| B-lineage cells (B220<sup>+</sup>) | 84.96 ± 3.326 N=5 | 75.27 ± 4.241 N=6 | NS |
| Pre-pro B cells (B220<sup>+</sup>/CD117<sup>+</sup>) | 2.183 ± 0.1974 N=6 | 2.567 ± 0.2951 N=6 | NS |
| Pro-B cells (IgM<sup>-</sup>/B220<sup>+</sup>/CD43<sup>high</sup>) | 7.380 ± 1.131 N=6 | 7.050 ± 0.9949 N=6 | NS |
| Pre-B cells (IgM<sup>-</sup>/B220<sup>+</sup>/CD43<sup>low</sup>) | 12.50 ± 1.965 N=6 | 13.54 ± 2.203 N=6 | NS |
| **Spleen** |  |  |  |
| B-lineage cells (B220<sup>+</sup>) | 58.16 ± 4.593 N=6 | 70.68 ± 3.524 N=6 | NS |
| Naïve mature B cells (B220<sup>+</sup>/IgM<sup>+</sup>/IgD<sup>+</sup>) | 45.09 ± 3.145 N=6 | 52.76 ± 1.874 N=5 | NS |
| Marginal zone B cells (B220<sup>+</sup>/CD21<sup>high</sup>/CD23<sup>low</sup>) | 7.235 ± 1.504 N=6 | 7.740 ± 0.5829 N=6 | NS |
| Follicular B cells (B220<sup>+</sup>/CD21<sup>low</sup>/CD23<sup>high</sup>) | 42.01 ± 3.506 N=6 | 45.75 ± 2.992 N=6 | NS |
| **Peritoneal cavity** |  |  |  |
| B-lineage cells (B220<sup>+</sup>) | 1.864 ± 0.3514 N=5 | 2.517 ± 0.2998 N=5 | NS |
| B1a cells (CD5<sup>+</sup>/IgM<sup>+</sup>) | 0.5924 ± 0.1002 N=5 | 0.8280 ± 0.1308 N=5 | NS |
| B1b cells (CD5<sup>-</sup>/IgM<sup>+</sup>) | 1.076 ± 0.2745 N=5 | 1.474 ± 0.2010 N=5 | NS |
| **Peyer's patches** |  |  |  |
| B-lineage cells (B220<sup>+</sup>) | 4.765 ± 1.106 N=6 | 5.841 ± 1.172 N=6 | NS |
| B220<sup>+</sup>/IgA<sup>+</sup> cells | 0.5625 ± 0.1273 N=6 | 0.8837 ± 0.2218 N=6 | NS |
| B220<sup>+</sup>/IgM<sup>+</sup> cells | 2.828 ± 0.5269 N=6 | 3.782 ± 0.8263 N=6 | NS |
| Naive B cells (B220<sup>+</sup>/PNA<sup>low</sup>/Fas<sup>low</sup>) | 3.684 ± 0.6369 N=5 | 3.556 ± 0.5026 N=5 | NS |
| Germinal centre B cells (B220<sup>+</sup>/PNA<sup>High</sup>/Fas<sup>High</sup>) | 1.044 ± 0.8822 N=8 | 3.258 ± 1.2 N=11 | P=0.0007 |

**Table S2.** SHM data (NGS) from individual mice in DNA repair proficient background. Total number of mutations, total number of bp analyzed and mutation frequencies for *w*t and *MARs*_*Eμ*_^Δ/Δ^ mice. (**A**) Data from intron 3’ to *J*_*H*_*4* in Peyer’s patches GC B cells, (**B**) Data from spleen GC B cells from SRBC-immunized mice, (**C**) Data from intron 5’ to Sμ in Peyer’s patches GC B cells.

| A | Intron 3' to J<sub>H</sub>4 Peyer's patch GC |  |  |  |  |  |
| --- | --- | --- | --- | --- | --- | --- |
|  | wt mice |  |  | MARSEμ<sup>Δ/Δ</sup> mice |  |  |
| # of individual mice | number of mutations | total number of bp analyzed | Frequency (mutation/Kb) | number of mutation | total number of bp analyzed | Frequency (mutation/Kb) |
| #1 | 5 919 179 | 434 026 896 | 13.6 | 1 835 118 | 193 099 971 | 9.5 |
| #2 | 1 381 154 | 127 267 013 | 10.9 | 3 217 763 | 388 192 563 | 8.3 |
| #3 | 7 063 941 | 539 404 010 | 13.1 | 1 224 927 | 230 604 186 | 5.3 |
| #4 | 2 213 918 | 148 184 470 | 14.9 | 4 215 306 | 354 052 562 | 11.9 |
| #5 | 4 027 597 | 378 109 299 | 10.7 | 1 974 487 | 210 307 731 | 9.4 |
| #6 | 1 402 298 | 123 363 043 | 11.4 | 1 429 132 | 250 627 288 | 5.7 |
| #7 | 2 498 967 | 182 824 193 | 13.7 |  |  |  |
| #8 | 2 105 343 | 208 395 146 | 10.1 |  |  |  |
| Total | 26 612 397 | 2 141 574 070 | 12.4 | 13 896 733 | 1 626 884 301 | 8.5 |

| B | Intron 3' to J<sub>H</sub>4 spleen GC (Immunized) |  |  |  |  |  |
| --- | --- | --- | --- | --- | --- | --- |
|  | wt mice |  |  | MARSEμ<sup>Δ/Δ</sup> mice |  |  |
| # of individual mice | number of mutations | total number of bp analyzed | Frequency (mutation/Kb) | number of mutation | total number of bp analyzed | Frequency (mutation/Kb) |
| #1 | 8 799 | 1 416 237 | 6.2 | 5 535 | 3 257 438 | 1.7 |
| #2 | 5 285 | 1 173 359 | 4.5 | 31 896 | 13 027 215 | 2.5 |
| #3 | 59 657 | 19 245 534 | 3.1 | 482 | 501 374 | 1.0 |
| Total | 73 741 | 21 835 130 | 3.4 | 37 913 | 16 786 027 | 2.3 |

| C | Intron 5' to SmuPeyer's patches GC |  |  |  |  |  |
| --- | --- | --- | --- | --- | --- | --- |
|  | wt mice |  |  | MARSEμ<sup>Δ/Δ</sup> mice |  |  |
| # of individual mice | number of mutations | total number of bp analyzed | Frequency (mutation/Kb) | number of mutation | total number of bp analyzed | Frequency (mutation/Kb) |
| #1 | 48604 | 35 947 796 | 1.4 | 340 669 | 70 101 915 | 4.9 |
| #2 | 42 937 | 32 953 275 | 1.3 | 1 098 398 | 215 265 958 | 5.1 |
| #3 | 59 309 | 28 156 682 | 2.1 | 530 553 | 94 311 231 | 5.6 |
| #4 | 17 594 | 11 681 044 | 1.5 | 241790 | 32530038 | 7.4 |
| #5 | 11 901 | 3231298 | 3.7 | 192 502 | 23871828 | 8.1 |
| #6 | 27 566 | 9864502 | 2.8 | 117 229 | 16868982 | 6.9 |
| #7 |  |  |  | 343 232 | 44846733 | 7.7 |
| Total | 207 911 | 121 834 597 | 1.7 | 2 864 373 | 497 796 685 | 5.8 |

**Table S3.** SHM data (NGS) from individual mice in genetic backgrounds deficient for base excision repair (*Ung*-deficient) and mismatch repair (*Msh2*-deficient) Total number of mutations, total number of bp analyzed and mutation frequencies. (**A**) Data from intron 3’ to *J*_*H*_*4* in Peyer’s patches GC B cells of *Ung* ^*Δ/Δ*^ *Msh2*^*Δ/Δ*^ and *Ung* ^*Δ/Δ*^ *Msh2*^*Δ/Δ*^ *MARs*_*Eμ*_^Δ/Δ^ mice, (**B**)Data from *Ung* ^*Δ/Δ*^ and *Ung* ^*Δ/Δ*^ *MARs*_*Eμ*_^Δ/Δ^ mice, (**C**) Data from intron 5’ to Sμ in Peyer’s patches GC B cells of *Ung* ^*Δ/Δ*^ *Msh2*^*Δ/Δ*^ and *Ung* ^*Δ/Δ*^ *Msh2*^*Δ/Δ*^ *MARs*_*Eμ*_^Δ/Δ^ mice, (**D**) Data from *Ung* ^*Δ/Δ*^ and *Ung* ^*Δ/Δ*^ *MARs*_*Eμ*_^Δ/Δ^ mice.

|  | Intron 3' to J<sub>H</sub>4 Peyer's patch GC |  |  |  |  |  |
| --- | --- | --- | --- | --- | --- | --- |
|  | UNG<sup>Δ/Δ</sup> MSH2<sup>Δ/Δ</sup> mice |  |  | UNG<sup>Δ/Δ</sup> MSH2<sup>Δ/Δ</sup> MARsEμ<sup>Δ/Δ</sup> |  |  |
| # of individual mice | number of mutations | total number of bp analyzed | Frequency (mutation/Kb) | number of mutation | total number of bp analyzed | Frequency (mutation/Kb) |
| #1 | 83 489 | 39 010 041 | **2.1** | 165 311 | 42 503 695 | **3.9** |
| #2 | 542 601 | 70 241 171 | **7.7** | 94 241 | 16 931 788 | **5.6** |
| #3 | 206 309 | 48 590 092 | **4.2** | 512 031 | 57 470 681 | **8.9** |
| #4 |  |  |  | 1 041 060 | 149 775 567 | **7** |
| #5 |  |  |  | 718 875 | 69 661 668 | **10.3** |
| Total | 832 399 | 157 841 304 | **5.3** | 2 531 518 | 336 343 399 | **7.5** |

**B**
|  | Intron 3' to J<sub>H</sub>4 Peyer's patch GC |  |  |  |  |  |
| --- | --- | --- | --- | --- | --- | --- |
|  | UNG<sup>Δ/Δ</sup> mice |  |  | UNG<sup>Δ/Δ</sup> MARsEμ<sup>Δ/Δ</sup> |  |  |
| # of individual mice | number of mutations | total number of bp analyzed | Frequency (mutation/Kb) | number of mutation | total number of bp analyzed | Frequency (mutation/Kb) |
| #1 | 272 454 | 26839331 | **10.2** | 613 978 | 89 213 320 | **6.9** |
| #2 | 1 798 561 | 145965068 | **12.3** | 473 338 | 44 306 794 | **10.7** |
| #3 | 432 270 | 160149949 | **2.7** | 441 712 | 67 366 607 | **6.6** |
| #4 | 633 350 | 90533829 | **7** | 569 070 | 65 436 018 | **8.7** |
| #5 | 507 031 | 47228276 | **10.7** | 817 177 | 73 685 315 | **11.1** |
| #6 | 99 600 | 9066959 | **10.9** | 744 638 | 79 286 350 | **9.4** |
| #7 | 208 684 | 19274286 | **10.8** |  |  |  |
| #8 | 258 678 | 34545602 | **7.5** |  |  |  |
| #9 | 318 557 | 30156482 | **10.6** |  |  |  |
| #10 | 1 576 407 | 163649903 | **9.6** |  |  |  |
| #11 | 2 822 217 | 195113384 | **14.5** |  |  |  |
| #12 | 563 458 | 56197886 | **10** |  |  |  |
| Total | 9 491 267 | 978 720 955 | **9.7** | 3 659 913 | 419 294 404 | **8.7** |

**C**
|  | Intron 5' to SmuPeyer's patches GC |  |  |  |  |  |
| --- | --- | --- | --- | --- | --- | --- |
|  | UNG<sup>Δ/Δ</sup> MSH2<sup>Δ/Δ</sup> mice |  |  | UNG<sup>Δ/Δ</sup> MSH2<sup>Δ/Δ</sup> MARsEμ<sup>Δ/Δ</sup> |  |  |
| # of individual mice | number of mutations | total number of bp analyzed | Frequency (mutation/Kb) | number of mutation | total number of bp analyzed | Frequency (mutation/Kb) |
| #1 | 78 467 | 34 863 085 | **2.3** | 64 593 | 11 768 040 | **5.5** |
| #2 | 117 529 | 19 114 688 | **6.2** | 794 845 | 81 456 430 | **9.8** |
| #3 | 4 285 | 698 559 | **6.1** | 829 374 | 83 165 361 | **10** |
| #4 |  |  |  | 394 268 | 34 269 808 | **11.5** |
| Total | 200 281 | 54 676 332 | **3.7** | 2 083 080 | 210 659 639 | **9.9** |

**D**
|  | Intron 5' to SmuPeyer's patches GC |  |  |  |  |  |
| --- | --- | --- | --- | --- | --- | --- |
|  | UNG<sup>Δ/Δ</sup> mice |  |  | UNG<sup>Δ/Δ</sup> MARsEμ<sup>Δ/Δ</sup> |  |  |
| # of individual mice | number of mutations | total number of bp analyzed | Frequency (mutation/Kb) | number of mutation | total number of bp analyzed | Frequency (mutation/Kb) |
| #1 | 154 039 | 46 862 890 | **3.3** | 566 494 | 78 527 619 | **7.2** |
| #2 | 138624 | 45 368 381 | **3** | 944 483 | 141 666 058 | **6.7** |
| #3 | 102 015 | 41 553 443 | **2.5** | 680 364 | 210 922 293 | **3.2** |
| #4 | 155 773 | 63 882 794 | **2.4** | 91 352 | 11 480 275 | **8** |
| #5 | 301 513 | 106 673 675 | **2.8** | 429 016 | 81 539 599 | **5.3** |
| #6 | 431 266 | 144 210 474 | **3** |  |  |  |
| #7 | 468 727 | 144 538 082 | **3.2** |  |  |  |
| Total | 1 751 957 | 593 089 739 | **3** | 2 711 709 | 524 135 844 | **5.2** |

